# Examining the Persistence of Human Coronaviruses on Fresh Produce

**DOI:** 10.1101/2020.11.16.385468

**Authors:** Madeleine Blondin-Brosseau, Jennifer Harlow, Tanushka Doctor, Neda Nasheri

## Abstract

Human coronaviruses (HCoVs) are mainly associated with respiratory infections. However, there is evidence that highly pathogenic HCoVs, including severe acute respiratory syndrome coronavirus 2 (SARS-CoV-2) and Middle East Respiratory Syndrome (MERS-CoV), infect the gastrointestinal (GI) tract and are shed in the fecal matter of the infected individuals. These observations have raised questions regarding the possibility of fecal-oral route as well as foodborne transmission of SARS-CoV-2 and MERS-CoV. Studies regarding the survival of HCoVs on inanimate surfaces demonstrate that these viruses can remain infectious for hours to days, however, to date, there is no data regarding the viral survival on fresh produce, which is usually consumed raw or with minimal heat processing. To address this knowledge gap, we examined the persistence of HCoV-229E, as a surrogate for highly pathogenic HCoVs, on the surface of commonly consumed fresh produce, including: apples, tomatoes and cucumbers. Herein, we demonstrated that viral infectivity declines within a few hours post-inoculation (p.i) on apples and tomatoes, and no infectious virus was detected at 24h p.i, while the virus persists in infectious form for 72h p.i on cucumbers. The stability of viral RNA was examined by droplet-digital RT-PCR (ddRT-PCR), and it was observed that there is no considerable reduction in viral RNA within 72h p.i.

## Introduction

Coronaviruses that infect humans (HCoV) belong to alpha and beta genera of the *coronaviridae* family. Four common HCoVs (229E, OC43, HKU1, and NL63) are responsible for 10-30% of common cold symptoms that can be mild to moderate *(18).* SARS-CoV-2, which is responsible for the COVID-19 pandemic, is a betacoronavirus that uses angiotensin conversion enzyme 2 (ACE-2) for entry. ACE-2 is abundantly expressed in the epithelium of the respiratory tract as well as the oral cavity, intestine and colon *(12, 20).* It is evident now that approximately 30-50% of COVID-19 patients demonstrate gastrointestinal symptoms including nausea, vomiting, diarrhea, and abdominal pain *(4, 21, 36).* SARS-CoV-2 RNA has been detected in more than 50% of patients’ stool specimens *(2, 11, 27, 30),* and several studies have confirmed that the virus detected in stool is infectious *(31, 37).* Moreover, persistent fecal viral shedding has been observed in pediatric patients *(33)* and there is direct evidence that SARS-CoV-2 can replicate productively in human enteroids and enterocytes *(12, 36).* More recently, it was demonstrated that multi-route mucosal inoculation (including oral inoculation) of African green monkeys with SARS-CoV-2 results in infection in both the respiratory and gastrointestinal tract *(10),* and orally inoculated golden Syrian hamsters develop respiratory and intestinal infection *(3).* Collectively, these observations suggest that fecal-oral transmission of SARS-CoV-2 is possible.

Although the primary route of transmission for HCoVs is inhalation of contaminated respiratory droplets and possible direct contact with contaminated fomites, there is concern that food could also act as a vehicle of transmission if contaminated with HCoVs. Food may become contaminated with HCoVs by contact with body secretions or fluids or by contact with soiled hands. Also, HCoVs may become aerosolized via talking, sneezing, or coughing of food-handlers and then be deposited on food surfaces. Food not only may act as a fomite, but can also transport the virus to the potentially susceptible oral cavity and the GI tract *(32).* There is evidence that certain HCoVs including HCoV-229E and MERS can survive GI conditions including low pH, digestive enzymes and bile *(38).* If this is the case for SARS-CoV-2, the relatively high viral titre in stool and rectal swabs of the infected individuals could be explained by active viral replication in the GI tract. Furthermore, fecal-oral is the main route of transmission for enteric coronaviruses such as swine coronaviruses *(26),* canine coronaviruses *(7),* and equine coronavirus *(19)* demonstrating that these viruses are not sensitive to the GI fluids.

Contamination of fresh produce may result in the transmission of not only the enteric viruses that are traditionally considered foodborne pathogens, but also possibly respiratory viruses such as adenoviruses, coronaviruses, and influenza viruses that can infect via contact with mucosal membranes {{640 O’Brien,B. 2020;}}. This is of particular concern for uncooked fruits and vegetables. Additionally, food handlers infected with respiratory viruses could still pose a potential health risk for food consumers, while preparing “cold foods” such as salads and sandwiches *(34).* Thus, it is imperative to examine the viral behaviour and inactivation in food and on food contact surfaces.

Since working with SARS-CoV-2 requires biosafety level 3 laboratory containment conditions, the use of surrogate HCoVs have been suggested to expand the current knowledge on coronavirus survival and inactivation under various conditions *(9).* For this reason, we chose HCoV-229E as a surrogate virus, since it has similar physicochemical properties to the more virulent HCoVs responsible for MERS and SARS *(29).* In this study, we examined the ability of HCoV-229E to retain infectivity on the surface of select fruits and vegetables, and thus obtained representative survival data that can be used to conduct risk assessments of SARS-CoV-2 transmission via food.

## Materials and Methods

### Cells and Viruses

HCoV-229E and human embryonic lung cell line MRC-5 were obtained from the American Type Culture Collection (CCL-171 and VR-740, respectively). Cells were grown at 37°C and 5% CO_2_ in culture media composed of Eagle’s minimal essential medium, supplemented with 0.23% (w/v) sodium bicarbonate, 500 μg/mL Penicillin-Streptomycin (ThermoFisher scientific), Glutamax-1, non-essential amino acids, and foetal bovine serum (FBS) 5% (v/v).

### Sample preparation

Three different produce types were tested: Royal Gala apples, Traditional Series tomatoes and English cucumbers (PLU code 4173, 4799 and 4593 respectively). Ten time points were selected, in triplicates: 0h, 0.5h, 1h, 2h, 4h, 6h, 16h, 24h, 48h and 72h. Each of the produce items was rinsed with water, dried with Kimwipes and disinfected with 70% ethanol. On the surface of each produce item, a 5cm by 5cm square was delimited using tape. This area was inoculated with 100μL of HCoV-229E (ATCC VR-740, 5×10^5^ PFU/mL). The liquid was spread using the tip of the pipette, then allowed to fully dry for 1h. After the appropriate time lapse at ambient conditions (22°C; relative humidity, 30% to 40%), the surface was sampled with a cotton swab, which was then placed into the MRC-5 culture media previously described *(17).* Samples were processed immediately after swabbing.

### Viral quantification

#### plaque assay

Viral quantification and survival time were determined by plaque assay using MRC-5 cells. Cells were grown at 37 °C and 5% CO_2_ in the culture medium previously described for up to three days, before being seeded, transferred into 12-well plates at a targeted concentration of 5×10^5^ cells/mL and incubated to reach a confluency of 80-90%. Samples were diluted in culture medium and 100μL of at least two dilutions were used in duplicate to infect the prepared plates for 90 min at 35°C and 5% CO_2_. Plates were manually rocked every 10 min during the infection phase. Cells were then washed with phosphate buffered saline (PBS) and covered with 2mL of overlay media, composed of a 50/50 mix of 2× culture medium previously described and 0.5% agarose. Plates were incubated at 35°C and 5% CO_2_ for 3-4 days. Cell monolayers were fixed using 3.7% paraformaldehyde for 4-24h, freed from overlay plugs by running under tap water and stained with 0.1% crystal violet for 20 min. Plaques were counted for each dilution to determine the viral titre.

#### Determining limit of detection

Each produce item was artificially inoculated with a serial dilution of the viral stock in triplicate. At T_0_, the virus was extracted and assayed by plaque assay as described above. The plaques were counted for each dilution and results were analyzed to determine the highest dilutions (lowest titre) for which plaques were still obtained in triplicate experiments.

#### Recovery rate calculation

The recovery efficiency was determined by calculating the ratio between the viral titre recovered at T_0_ and the viral titre that was used to inoculate the sample.

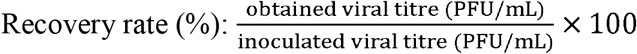

#### Estimating the decay rate

Viral decay rate was calculated as described previously *(13).* Briefly, linear regressions of the natural logarithm of virus abundance versus time (in hours) was calculated. The slope of the regressions represent the decay rate and when multiplied by 100, represent percentage of infectivity lost per hour. Viral half-life was calculated by dividing ln(2) by the slope.

#### ddRT-PCR

For each produce item, all triplicates of 10 time points were tested. Viral RNA was isolated using a QIAamp viral RNA kit (QIAGEN) and diluted in sterile molecular biology grade water (Corning). The QX200 ddPCR system (Bio-Rad) was used for quantification and all PCR reactions were prepared using the One-Step RT-ddPCR Advanced Kit for Probes (Bio-Rad Cat# 1864022). Primers used were previously described in *(25):* Forward primer 229E-FP (5-TTCCGACGTGCTCGAACTTT-3; GenBank accession no. M33560; nt 474 to 493) and reverse primer 229E-RP (5-CCAACACGGTTGTGACAGTGA-3; nt 523 to 543). A new probe that would complement the primers and be compatible with TaqMan qPCR requirements (ABI 7700 Users Manual) was designed by using Integrated DNA Technologies (IDT) OligoAnalyzer tool. The new probe had the appropriate dissociation temperature and a minimal likelihood for duplex or hairpin formation: 229E-PR (5’-/56-FAM/TGCATTGAC/ZEN/CTCAGGATTCCATGCCC/3IABkFQ/-3’). Each PCR reaction contained 5μL of RNA, 1000 nmol/L of each primer, and 280 nmol/L of each probe. All samples were tested in duplicate. Droplets were generated using the QX200 droplet generator (Bio-Rad) according to the manufacturer’s protocols, and PCR was performed using the following cycling conditions: an initial reverse transcription at 48°C for 30 min, followed by PCR activation at 95°C for 10 min and 45 cycles of amplification (15 s at 95°C and 1 min at 60°C). Droplets were detected in the QX200 droplet reader and analyzed using the Quantasoft version 1.7.4.0917 (Bio-Rad) software.

## Results

### Recovery Efficiency from Produce

As shown in Table 1, the recovery efficiency of HCoV-229E from all the tested commodities is well above 1%, with the highest recovery rate (10.8%) from tomatoes and the lowest (4.1%) from cucumbers. The limit of detection (LOD) for each commodity is determined as the lowest spiking concentration that produced plaques for all three replicates. As indicated in Table 2, the LOD was approximately 125 PFU for tomatoes and apples, and 50 PFU for cucumbers.

**Table 1.** Recovered viral titre at T_0_ and recovery rate in percentage for each produce type. The results are the mean of 3 independent experiments.

| Produce | Titer at T<sub>0</sub><br>(PFU/mL) | Recovery rate<br>(%) |
| --- | --- | --- |
| Apple | 1.45E+03 | 5.81 |
| Tomato | 2.69E+03 | 10.77 |
| Cucumber | 1.20E+03 | 4.09 |

**Table 2.** Detection of HCoV-229E on the surface of different produce. Samples were inoculated with 10^4^ to 10^1^ PFU of HCoV-229E and examined by plaque assay at T_0_. ND is not detected.

| Produce | Viral Inoculum (PFU) |  |  |  |  |  |  |
| --- | --- | --- | --- | --- | --- | --- | --- |
|  | 10,000 | 1000 | 500 | 250 | 125 | 50 | 10 |
| Apple | 3/3 | 3/3 | 3/3 | 3/3 | 3/3 | ND | ND |
| Tomato | 3/3 | 3/3 | 3/3 | 3/3 | 3/3 | 2/3 | ND |
| Cucumber | 3/3 | 3/3 | 3/3 | 3/3 | 3/3 | 3/3 | ND |

### Persistence of infectivity

We artificially inoculated the surface of apples, tomatoes and cucumbers with 5×10^4^ PFU of HCoV-229E, which is consistent with the amount of virus that is typically exhaled by an infected individual *(14).* Figure 1 shows the persistence in infectivity of HCoV-229E at RT within 72 h p.i. The change in infectious viral titre is similar in apples and tomatoes with a progressive decline in infectivity up to 16h p.i. (Figure 1, Table 3). No infectious viral particles were isolated from tomatoes and apples at 24 h p.i., which demonstrates that viral infectivity is reduced below the LOD (i.e. >3 log reduction). However, infectious viral particles were detected on cucumbers up to 72 h p.i. Within the first 4 h p.i, viral infectivity reduces over 1 log on tomatoes and apples (1.18 and 1.27 log, respectively), while the reduction on cucumbers is only 0.75 log (Table 3). The reduction in infectivity is less than 2 log at 24 h p.i on cucumbers and by 72 h p.i. reaches approximately 2.5 log. No infectious viral particles were detected on cucumbers at 96 h p.i.

**Figure 1.**
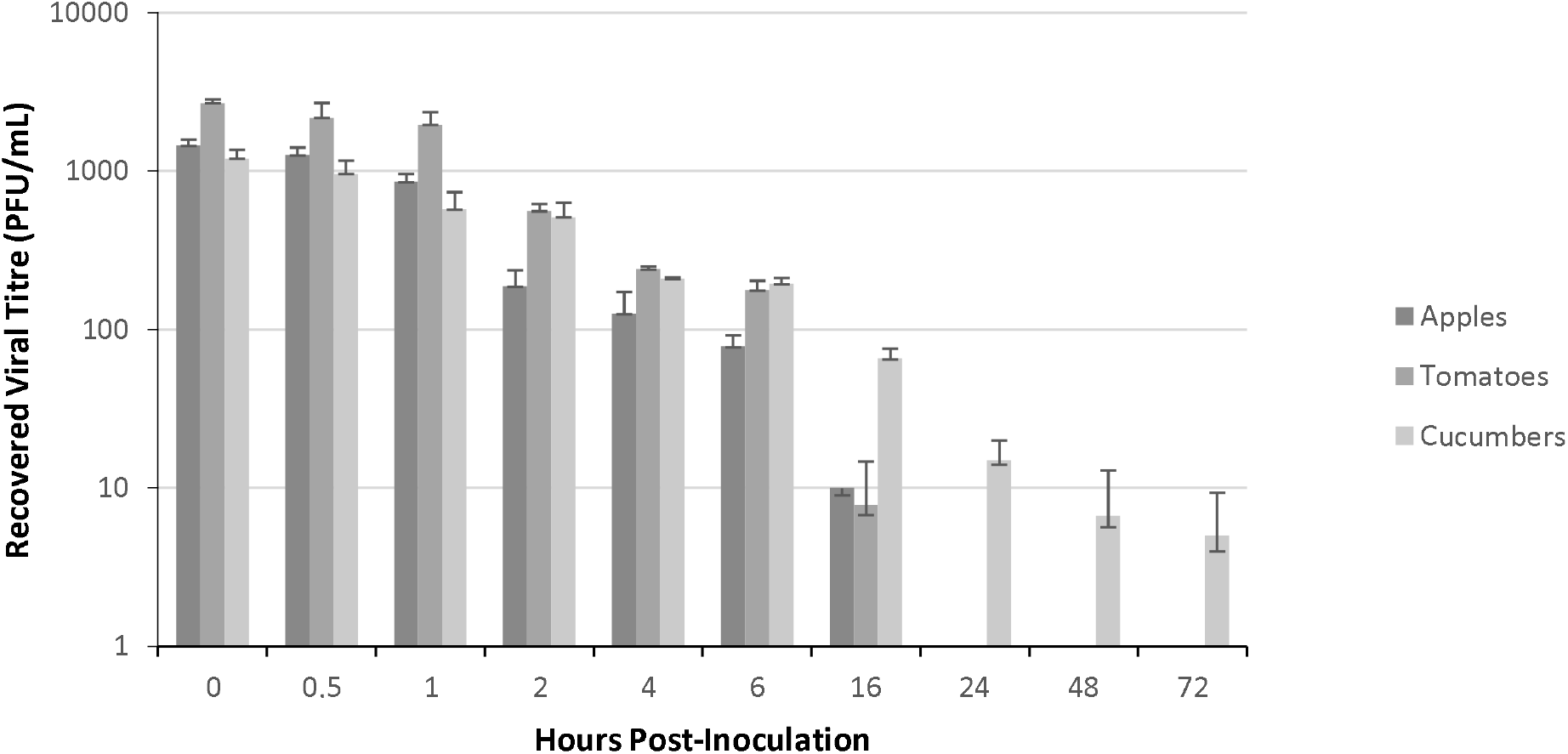
Persistence of infectious HcoV-229E on commonly consumed fruits and vegetables. Approximately 5×10^4^ PFU HCoV-229E (100 μl viral stock) was applied to the tested surface and incubated at ambient conditions (22°C; relative humidity, 30% to 40%). Virus was extracted and assayed for infectivity at various time points as described in the text. The data represent the average of three independent experiments. Error bars represent standard deviation.

**Table 3.**
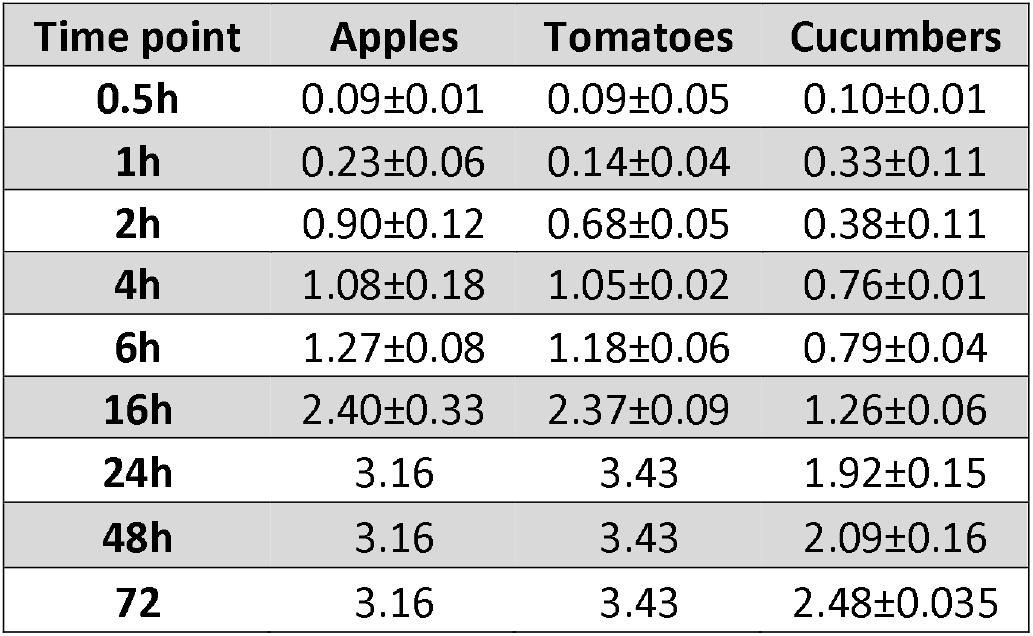
Log reduction in viral titre compared to T_0_. The results are the mean of 3 independent experiments ± Standard Deviation.

The median decay rate of HCoV-229E on apples and tomatoes was similar at 30%/h and 34%/h respectively, while the median decay rate on cucumbers was considerably lower at 7.7%/h. The median half-life of the virus on apples and tomatoes was 2.3h and 2.05h respectively and the median half-life on cucumbers was 9.05h (Table 4).

**Table 4.** Decay rate (DR) in percentage and viral half-life (HL) in hours (h) on each produce type. The results are the median of 3 independent experiments ± Standard Deviation.

|  | DR (%) | HL (h) |
| --- | --- | --- |
| **Apple** | 30±0.25 | 2.3±0.02 |
| **Tomato** | 34±0.1 | 2.05±0.06 |
| **Cucumber** | 7.7±0.6 | 9.05±0.75 |

### Persistence of viral RNA

We next set out to investigate the persistence of viral RNA on the examined produce over 72 h.p.i. at ambient temperature. As demonstrated in Figure 2, no drastic reduction in viral RNA titre was observed over a 72h p.i. period. On apples, tomatoes, and cucumbers, viral RNA decreased by approximately 0.7 log, 0.5 log, and 0.3 log, respectively compared to T_0_,. Altogether, these observations demonstrate that viral RNA is more resistant to degradation compared to viral infectivity on the surface of produce.

**Figure 2.**
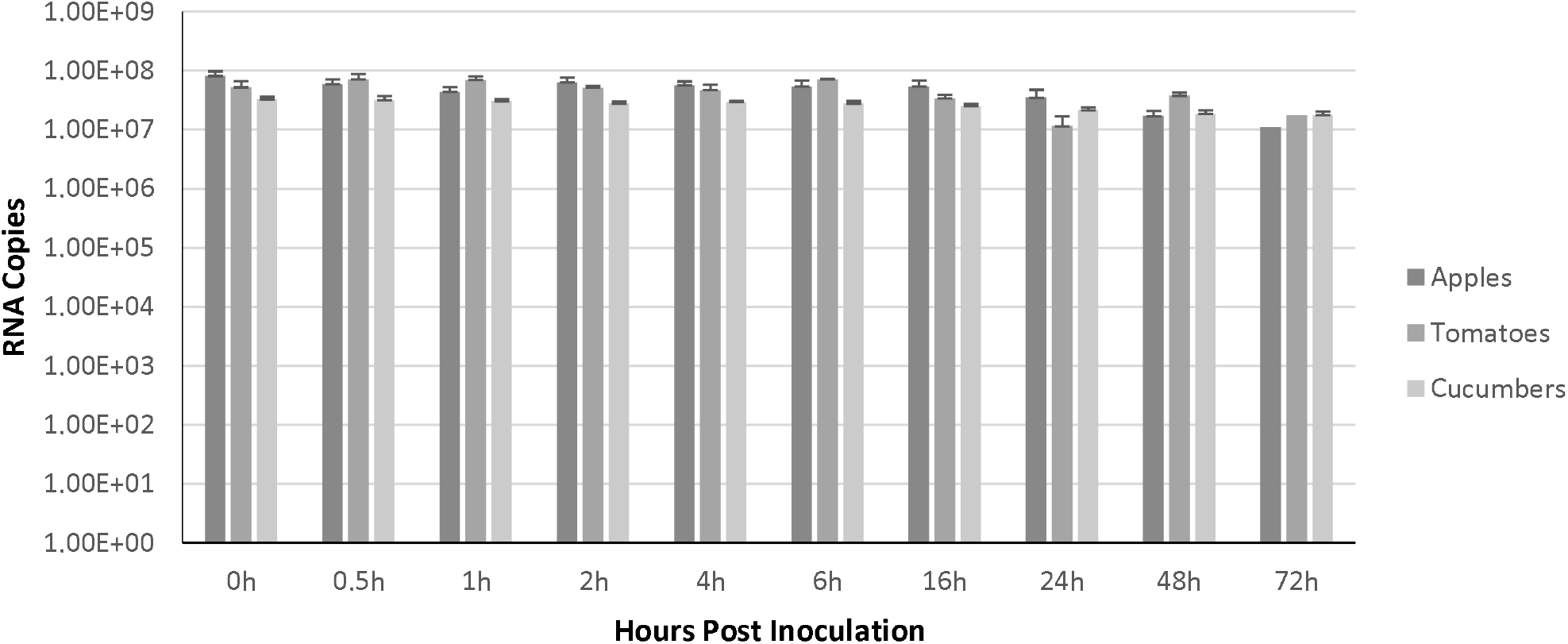
Persistence of viral RNA on commonly consumed fruits and vegetables. Approximately 2×10^8^ RNA copies of HCoV-229E (100 μl of viral stock) was applied to the tested surface and incubated at ambient conditions (22°C; relative humidity, 30% to 40%). Virus was extracted at indicated time points and viral RNA was quantified by ddRT-PCR. The data represent the average of three independent experiments. Error bars represent standard deviation.

## Discussion

To date, there is no conclusive evidence of foodborne transmission of SARS-CoV-2, however, the traditional epidemiological foodborne investigation is unlikely to be employed with COVID-19 patients. For example, it is unlikely that infected people are asked to recall foods that they may have consumed during the period when they became infected. Without this information, any association between SARS-CoV-2 and foods cannot be made, and understanding the role of foodborne transmission remains elusive. Obtaining this epidemiological information would be helpful for efficient contact-tracing and source-tracking as more than 54% of COVID-19 patients can not recall how and where they contracted the virus *(23).*

Environmental persistence of HCoVs has been examined by different groups, who have obtained contradictory results *(1).* One study has shown that the stability of SARS-CoV-2 and SARS-CoV-1 on dry surfaces at RT is similar, with no infectious virus being retrieved after 72h p.i. *(24),* while, Chin et al recovered infectious SARS-CoV-2 from plastic and stainless steel up to 7 days p.i. *(5).* Keevil and coworkers reported that HCoV-229E remains infectious for 5 days at RT on a range of surface materials including glass and PVC, while it is rapidly inactivated on the surface of copper alloys *(28).* In another study, more relevant to this work, it was shown that the infectivity of HCoV-229E is completely abolished within 4 days p.i. on lettuce at 4°C *(34).* Recently, it was demonstrated that SARS-CoV-2 remains infectious on salmon at RT for 2 days *(15).* Herein, we only examined viral survival at ambient temperature and we have shown the infectivity of HCoV-229E is reduced to below LOD followed by 24h incubation on tomatoes and apples, and 96h on cucumbers.

At this point, we speculate that the longer survival on cucumbers compared to apples and tomatoes could be partly explained by the difference in surface pH of these commodities. The influence of pH on the stability of several coronaviruses has been studied and it has been shown that in general, coronaviruses are more stable at near neutral pH as compared to acidic or alkaline pH *(1).* As such, the near neutral surface pH of cucumbers (5.7), compared to the more acidic surface pH of tomatoes and apples (4.2 and 3.9, respectively), could be more suitable for the survival of HCoV-229E *(16).* It should also be noted that the LOD on cucumbers was lower compared to apples and tomatoes (50 PFU compared with 125 PFUs, respectively). Thus, it is possible that HCoV-229E remained infectious by 24 h p.i. on apples and tomatoes but the titre was below the LOD. However, the decay rate on cucumbers is considerably slower compared to apples and tomatoes (Figure 1 and Table 4), and the viral half-life on cucumbers is very close to the viral half-life on plastic *(24)* (9.05h and 9.04h, respectively). Further investigation is needed to determine whether the surface of apples and tomatoes has some virucidal properties, not found on inanimate surfaces, that may lead to a more rapid viral inactivation. Thus, our results are in accordance with the previous findings that HCoVs lose their infectivity within a few days on inanimate surfaces at RT *(22).* Therefore, if produce becomes contaminated with HCoVs through irrigation or contaminated hands during pre- or post-harvest, while being stored at ambient temperature, the risk will be considerably reduced by the time it reaches the consumers. However, if the contamination occurs at the end of the food processing chain, for example by infected personnel in a restaurant setting, where the prepared food is consumed within a few minutes, there is a potential risk for infection. In such scenarios, the risk of super-spreading events is high as well *(6, 35).*

The persistence of viral RNA on the studied produce for several days despite the loss of infectivity, can be explained by the high environmental resilience of the coronavirus shell, which protects the viral genome *(8).*

It should be noted that our study involved experimental inoculation of fresh produce with HCoV-229E, and thus may not be fully representative of potential natural contamination. However, the infectious titre of virus used for inoculation of samples in the current study is representative of a worst-case scenario, if virus was found to be present on fresh produce. Herein, we attempted to address an important knowledge gap regarding the survival of human coronaviruses on fresh produce at ambient temperature. Potential foodborne transmission poses important public health implications and may partly explain the possible recurrence of the disease and its persistent transmission. Thus, our results could support more robust decisionmaking concerning risk assessment for foodborne transmission of human coronaviruses.

## Acknowledgements

The authors would like to thank Dr. Brent Dixon and Dr. Franco Pagotto from the Bureau of Microbial Hazards for kindly reviewing the manuscript and providing insightful comments.

